# ELAVL1 Primarily Couples mRNA Stability with the 3’UTRs of Interferon Stimulated Genes

**DOI:** 10.1101/2020.08.24.263418

**Authors:** Katherine Rothamel, Sarah Arcos, Byungil Kim, Clara Reasoner, Neelanjan Mukherjee, Manuel Ascano

**Affiliations:** Department of Biochemistry, Vanderbilt University School of Medicine, Nashville, TN, 37232, USA; Department of Biochemistry and Molecular Genetics, University of Colorado School of Medicine, Aurora, CO, 80045, USA

**Keywords:** innate immunity, ELAVL1, binding site, 3’UTR, RNA stability

## Abstract

Upon detection of a pathogen, the innate immune system triggers signaling events leading to the transcription of mRNAs that encode for pro-inflammatory and anti-microbial effectors. RNA-binding proteins (RBPs) interact with these functionally critical mRNAs and temporally regulate their fates at the post-transcriptional level. One such RBP is ELAVL1, which is known to bind to introns and 3’UTRs. While significant progress has been made in understanding how ELAVL1 regulates mRNAs, how its target repertoire and binding affinity changes within an immunological context remains poorly understood. Here, we overlap four distinct high-throughput approaches to define its cell-type and context-dependent targets and determine its regulatory impact during immune activation. ELAVL1 overwhelmingly binds to intronic sites in a naïve state, but during an innate immune response, ELAVL1 targets the 3’UTR - binding both previously and newly expressed mRNAs. We find that ELAVL1 mediates the RNA stability of genes that regulate the pathways involved in pathogen sensing and cytokine production. Our findings reveal the importance of examining RBP regulatory impact under dynamic transcriptomic events to best understand their post-transcriptional regulatory roles within specific biological circuitries.

## INTRODUCTION

The need to rapidly control gene expression is of paramount importance to implement a robust but punctuated immune response. When a cell is exposed to an immunogenic stimulus, high levels of interferon-stimulated gene (ISG) transcripts are expressed, requiring the cell to orchestrate their translation while simultaneously preventing pathogenic (e.g., viral) RNA from using the same machinery (Liu and Qian, 2013; Piccirillo et al., 2014). ISGs encode for anti-viral, pro-inflammatory, and survival proteins and their expression is essential in creating a heightened immunoreactive state (Hubel et al., 2019; Schneider et al., 2014). Just as essential as the initiation of an inflammatory response are the processes that lead to its resolution; therefore, the cell must return to a basal state by limiting the activities of ISGs to prevent damage to the host tissue (Anderson, 2009; Khabar and H. A. Young, 2007; Rigby and Rehwinkel, 2015; Savan, 2014). Prolonged Interferon-Beta 1 (IFNB1) expression has been shown to increase susceptibility to many inflammatory diseases and is a hallmark of autoimmune diseases and cancer (Crow, 2015; Frangou et al., 2013; Reder, 2013). Emerging evidence indicates that RNA-binding proteins (RBPs) can affect the levels and translation rates of immune specific transcripts to influence the intensity and duration of an innate immune response (Anderson, 2008; Hao and Baltimore, 2009; Kafasla et al., 2014; Mino and Takeuchi, 2013). RBPs facilitate RNA metabolism through the control of such processing events as splicing, subcellular localization, stability, and translation (Dreyfuss et al., 2002; Gerstberger et al., 2014; Keene, 2007; Lunde et al., 2007). RBPs act *in trans* by binding specific structural and/or sequence *cis*-elements, often within the 3’UTR of mRNAs, a highly trafficked region that is essential to many modes of post-transcriptional gene regulation (Gebauer et al., 2012).

A major strategy by which RBPs function to regulate mRNAs during immune-activated states is through managing their stability. RBPs such as Tristetraprolin (TTP) and the cytotoxic granule-associated T-Cell-Restricted Intracellular Antigen 1 (TIA1) function as negative regulators of cytokines by leading to increased transcript decay, which is essential for the resolution of inflammation (Herman et al., 2018; Meyer et al., 2018; Tiedje et al., 2016). Conversely, Embryonic Lethal Vision-Like Protein 1 (ELAVL1) has been reported to play a role in immunoregulation by antagonizing the effects of RBPs such as TTP (Kafasla et al., 2014). ELAVL1, also known as HuR (Szabo et al., 1991), binds uridine-(U) and adenyl-uridine-rich elements (AREs) (Chen and Shyu, 1995; López de Silanes et al., 2004), a common low complexity *cis*-element found throughout the transcriptome. Yet, somewhat paradoxically, ELAVL1 binds almost exclusively to cellular mRNAs – more specifically at introns and 3’UTRs (Lebedeva et al., 2011; Mukherjee et al., 2011; Sedlyarov et al., 2016). ELAVL1 is ubiquitously expressed in most cell types and has three distinct and highly conserved RNA-binding domains belonging to the RNA-recognition motif (RRM) family. During steady-state conditions, ELAVL1 is predominantly found in the nucleus but can translocate to the cytoplasm via phosphorylation of Y200, S202, and S221, located in the hinge region of the protein between the second and third RRM. The phosphorylation of ELAVL1 at these residues is reported to occur as part of signal transduction events, including cellular response to immune agents and mitogen signal transduction events (X. C. Fan and Steitz, 1998; Grammatikakis et al., 2016).

Previous reports have shown that ELAVL1 is required to maintain the mRNA levels of AU-containing transcripts, including the immune relevant transcripts *IFNB1, COX-2, IL-8*, and *TGFB* (Brennan and Steitz, 2001; Dixon et al., 2001; J. Fan et al., 2011; X. C. Fan and Steitz, 1998; Herdy et al., 2015; Peng et al., 1998). However, most of these studies do not examine if these effects on mRNA levels are due to the direct binding of ELAVL1. Additionally, many of these studies examine the regulatory impact of ELAVL1 on a singular target. Thus, how ELAVL1 prioritizes cellular targets and orchestrates its role in overall immunoregulation was not fully ascertained. Further complicating our understanding of the role of ELAVL1 in immunity and inflammation are the phenotypic outcomes reported in mouse models (Christodoulou-Vafeiadou et al., 2018; Katsanou et al., 2005; Srikantan et al., 2012). ELAVL1 knockouts in murine cells have conflicting results depending on the cell-type (epithelial vs. myeloid) and opposing phenotypes depending on the pattern recognition receptor (PRR) agonist (LPS vs. RIG-I) (Yiakouvaki et al., 2012).

With the advent of high-throughput sophisticated RBP-crosslinking and immunoprecipitation (CLIP) methods, such as PhotoActivable Ribonucleoside-enhanced Cross-Linking and Immunoprecipitation (PAR-CLIP), the ability to precisely capture the binding sites of RBPs such as ELAVL1 in cells and study their direct effects have enabled a more molecular understanding of their function (Hafner et al., 2010). The targets and the regulatory impact of ELAVL1 on mRNA targets have been performed in HEK293 and HeLa cells (Lebedeva et al., 2011; Mukherjee et al., 2014). These landmark studies ushered in a broader appreciation for the post-transcriptional role that ELAVL1 can elicit on its targets – particularly for pre-mRNA processing. However, these earlier reports examined ELAVL1 regulation of targets under steady-state conditions in cell types that do not reflect more specific biological processes for which ELAVL1 is implicated - making it difficult to extrapolate whether the reported direct targets contribute to the phenotypes associated with overexpression or knockout of ELAVL1 in murine models. This is especially true for an RBP that binds to commonly occurring ARE’s – which obviates the effectiveness of using predictive analyses to identify its functional targets. Given its purported biological roles in development, cancers, and immunoregulation, what remains lacking is a deeper understanding of how the targeting and binding affinity of ELAVL1 to RNA changes in response to a cellular signaling event that substantively alters the transcriptome, and by virtue, substrate pool of ELAVL1.

Here, we report a multidisciplinary analysis of the targeting and functional outcomes of ELAVL1 during a nucleic acid-induced innate immune response in human THP-1 monocyte-like cells. We find that the mRNA targets of ELAVL1 in THP-1 cells only share 25% of the binding sites of previously published datasets. ELAVL1 largely transitions to binding the 3’UTRs of mRNA transcripts upon innate immune activation, and this 3’UTR shift is an absolute prerequisite for enrichment. The loss of ELAVL1 led to widespread destabilization of its enriched target transcripts. Specifically, we found that highly regulated targets had a three-fold average reduction in their stabilities, losing 30 to 80% of their original half-lives. Importantly, we found that among the most highly regulated targets were transcripts that encode for ISGs and their transcriptional regulators, suggesting that ELAVL1 contributes at multiple levels of a pro-inflammatory response. To date, this is the first report comparing how the binding properties and the targeting of an RBP change between steady-state and innate immune conditions, thus providing a general framework for investigating RBPs as they govern dynamic transcriptomes.

## RESULTS

### Transcriptional landscape during an IRF3 innate immune response

To model a nucleic acid-induced innate immune transcriptional response, which would be analogous to viral infection and cellular detection of pathogenic nucleic acids, we stimulated THP-1 monocyte-like cells with cyclic GMP-AMP (cGAMP) - the endogenous agonist of the immune adaptor protein STIMULATOR OF INTERFERON GENES (STING) (Cai et al., 2014; Ishikawa and Barber, 2008; Sun et al., 2013; Wu et al., 2013). cGAMP is a second messenger molecule produced by the pattern recognition receptor cyclic GMP-AMP synthase upon the detection of cytoplasmic dsDNA in the cytoplasm (Gao et al., 2013a; 2013b). cGAMP activation of STING ultimately elicits an INTERFERON REGULATORY FACTOR-3 (IRF3)-dependent transcriptional response, which includes the upregulation of hundreds of ISGs (Cai et al., 2014; Ishikawa and Barber, 2008; Ishikawa et al., 2009; Sato et al., 1998; Shae et al., 2019). Importantly, the activation of IRF3 integrates not only the detection of cytoplasmic dsDNA but also the upstream activities of the RIG-I family of RNA pattern recognition receptors (Honda et al., 2006). Thus, exposing THP-1 cells to cGAMP allows for precise and robust activation of cells via IRF3 – a major arm of the innate immune system, without confounding crosstalk effects through the stimulation of additional immune signaling pathways by other pathogen-associated molecular pattern molecules.

Against the transcriptomic background of an IRF3-driven innate immune response, we sought to characterize the post-transcriptional gene regulatory role of ELAVL1. As an overall experimental design, we integrated four independent high-throughput datasets (Figure 1A). Using RNA-Sequencing (RNA-Seq), we (i) performed gene expression analysis comparing the mRNA levels from naïve and cGAMP-stimulated THP-1 cells. We next (ii) identified the direct RNA targets of ELAVL1 in both cellular conditions using PAR-CLIP (Hafner et al., 2010). Using RNA Immunoprecipitation (RIP)-Sequencing, we (iii) quantified the relative enrichments of ELAVL1 targets (Keene et al., 2006; Ramanathan et al., 2019; Tenenbaum et al., 2000; Zhao et al., 2010). Finally, we (iv) assessed the regulatory impact of ELAVL1 by measuring the RNA half-lives of its target mRNAs with thiol (SH)-Linked Alkylation for Metabolic Sequencing of RNA (SLAM-Seq) upon the loss of its expression during an immune-stimulated transcriptional state (Herzog et al., 2017). Together, these datasets will enable the identification of high-confidence ELAVL1-regulated targets during an innate immune response and provide insight on how signaling events alter the mRNA targets of an RBP, like ELAVL1, during a changing substrate landscape.

**Figure 1.**
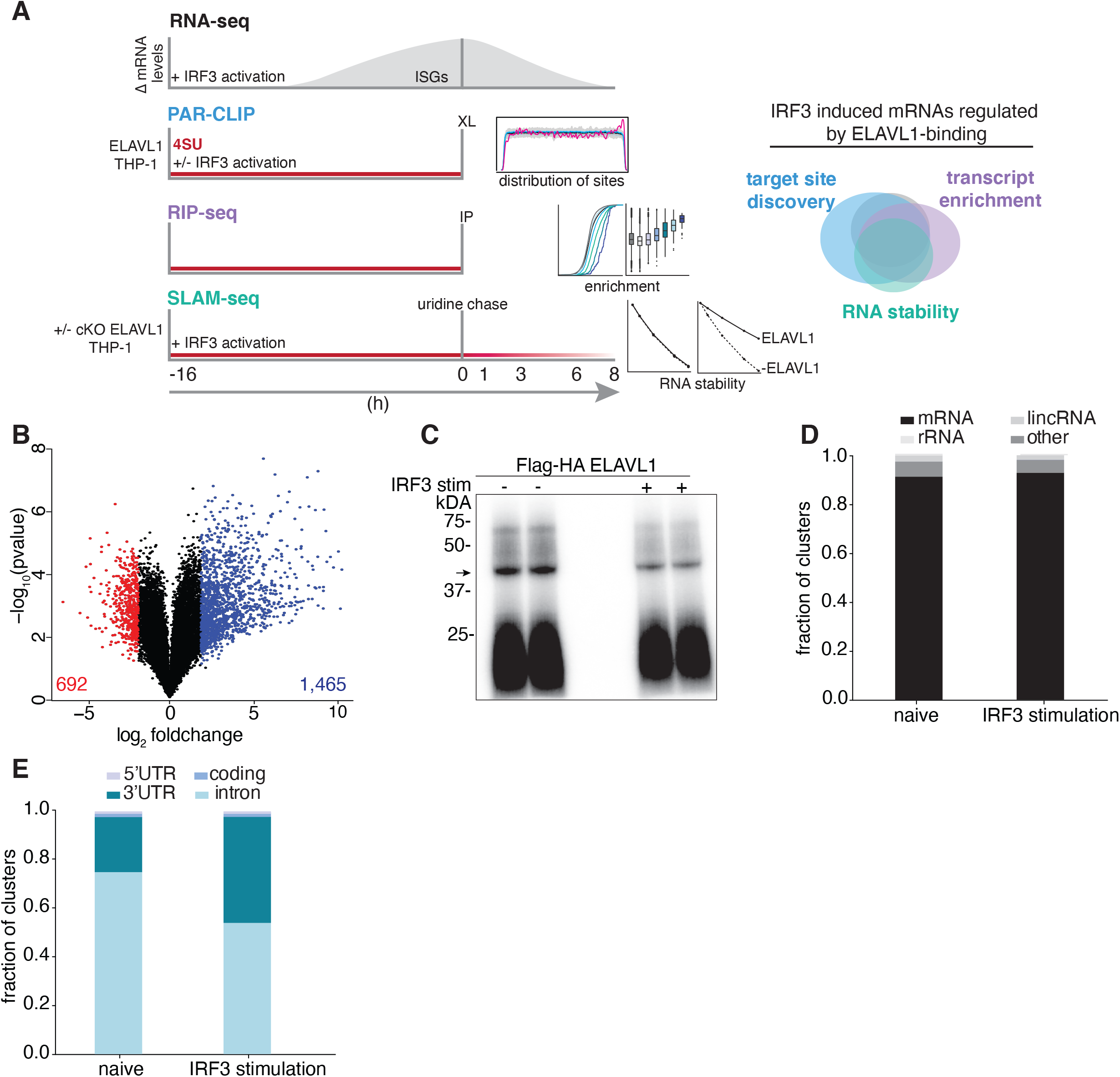
RNA-Seq and PAR-CLIP capture the context-dependent RNA substrates of ELAVL1. (**A**) Schematic of the overall experimental design used to define how ELAVL1 regulates its mRNA substrates during an innate immune response. In naïve and stimulated THP-1 cells, mRNA levels of potential ELAVL1 substrates were measured using RNA-Seq. Then condition-specific ELAVL1 binding site data were mapped at nucleotide resolution to RNA loci using PAR-CLIP. RIP-seq was used to quantitatively measure the interaction ELAVL1 has to its RNA targets. Lastly, ELAVL1 target stability was measured using SLAM-seq. Venn diagram (right panel) illustrates how high throughput datasets will be used to assess the functional targets of ELAVL1. (**B**) Volcano plot comparing the differential mRNA levels between naïve and stimulated THP-1 cells. The red and blue represent the down- and up-regulated transcripts in response to cGAMP stimulation. (**C**) Phosphorimage of ^32^P-RNA bound to ELAVL1 in the naïve and stimulated THP-1 cells. 4SU crosslinked ELAVL1 was immunoprecipitated then separated by SDS-PAGE; arrow indicates ELAVL1 and covalently-bound and radiolabeled RNAs. (**D**) Distribution of ELAVL1-binding sites across indicated RNA categories. (**E**) Distribution of binding sites across different mRNA transcript features for naïve and stimulated PAR-CLIP samples.

To compare the mRNA levels that occur in THP-1 cells upon 16 h exposure to cGAMP, we performed gene expression profiling using RNA-Seq (Figure 1B). The 16 h time point was selected based on our assessment of peak level expression of ISGs, including IFNB1, which was previously identified as a direct target of ELAVL1 (Figure S1A) (Herdy et al., 2015; Takeuchi, 2015a). We found that 2,157 (1,465 upregulated and 692 downregulated) genes were differentially regulated with two-fold or greater change (adjusted *P*-value ≤ .005) upon stimulation (Figure 1B, Table S1). Pathway analysis of the top (25%) upregulated genes indicated that these mRNAs are involved in the regulation of the innate immune response, apoptosis, and hematopoiesis (not shown). Many of the upregulated genes (60%) are known ISGs including the anti-viral effectors (*OAS, MX1*, *ISG15*), positive regulators of IFN response (*IRFs, STAT1, JAK*), and nucleic acid pattern recognition receptors (*TLR8, RIG-I, IFITs*) (Hubel et al., 2019; Rusinova et al., 2012; Schneider et al., 2014; Schoggins et al., 2011).

### The binding preference and mRNA targets of ELAVL1 are cell-type and context-dependent

To identify the direct RNA-targets of ELAVL1 during a naïve-state (naïve) and an IRF3-driven innate immune state (stimulated), we performed PAR-CLIP in THP-1 cells conditionally expressing Flag and HA epitope-tagged ELAVL1 (Flag-HA ELAVL1) that were either stimulated with cGAMP for 16 h or mock-treated prior to crosslinking. A phosphorimage of the crosslinked and immunoprecipitated Flag-HA ELAVL1 revealed one major band migrating at approximately 40 kDa, the expected molecular mass of Flag-HA ELAVL1, in the presence and absence of an immune stimulus (Figure 1C). RNA from this band was recovered and processed for small RNA-Seq. Each cDNA library contained approximately 70 million reads with an average of ~10.7 and ~12.7 million *unique* reads for the naïve and stimulated samples, respectively (Table S2). ~91% of the reads mapped to (pre-)mRNA with an 86.4% average T-to-C fraction across all samples from naïve and stimulated conditions, altogether indicating high-quality recovery and crosslinking efficiencies of our PAR-CLIP procedure for isolating ELAVL1-bound RNA targets (Figure 1D). Using PARpipe, we identified 133,740 naive, and 50,074 stimulated ELAVL1 distinct RNA-binding sites that have ≥ 2 unique T-to-C positions and > 20% T-to-C ratio (Corcoran et al., 2011). Although the greater than two-fold difference in the total number of RNA-binding sites between the two conditions was notable, this difference was not due to lower complexity libraries for the stimulated samples since these libraries contained more unique reads mapped per cluster (Figure S1B). 106,081 (79%) and 40,681 (81%) of the clusters identified by PAR-pipe have RNA-Seq expression data (> 5 counts per million, CPM) and RIP-Seq enrichment data (IgG normalized) for the RNA transcripts that correspond with the cluster (Table S3). Of those clusters, 98,689 (74%) and 38,607 (76%) of ELAVL1 binding sites map to (pre)-mRNA with high confidence in naïve and stimulated samples, respectively.

To test the hypothesis that ELAVL1 has distinct binding-preferences due to cell-type, we compared the THP-1 naive binding sites to previously published PAR-CLIP data on ELAVL1 from HEK293 (Mukherjee et al., 2011). 29,820 clusters (30% of the THP-1 naïve clusters and 25% of the HEK293 clusters) overlapped by at least one nucleotide between the THP-1 and HEK293 datasets (Figure S1C). We find that ELAVL1 in naïve THP-1 cells uniquely binds to 1,725 mRNAs even though 1,007 (58%) of these transcripts are expressed in HEK293 based on RNA-Seq data (Mukherjee et al., 2011). Reactome pathway analysis revealed that the mRNAs bound by ELAVL1 in THP-1 cells are associated with NF-kB activation (R-HSA-933543), viral defense (R-HSA-168273), and toll-like receptor signaling (R-HSA-5603041) (Figure S1D).

We found that a substantial fraction of ELAVL1 binding sites (74%) in THP-1 cells were located within the introns of pre-mRNAs in the naïve state, as similarly observed in HEK293 cells (Mukherjee et al., 2011). However, upon immune stimulation, intronic binding was significantly depleted: only 52% of the clusters mapped to introns of target transcripts in the stimulated condition (Figure 1E, Table S2). Nonetheless, intron-located clusters were enriched near 5’ and 3’ splice sites in both conditions suggesting a consistent role for ELAVL1 in regulating splicing (Figure S1E). Nearly 96% of the exonic binding sites mapped to the 3’UTR in both naïve and immune activated conditions. The 3’UTR-binding sites were depleted proximal to the stop codon (0-150 nts after stop codon) and enriched toward the most distal region of the transcript near the polyadenylation site (40 nts before the poly-A site) (Figure S1F and S1G). Interestingly, when we compared the distribution of the binding sites along the 3’UTR to previous ELAVL1 PAR-CLIPs (Lebedeva et al., 2011), we found that the enrichment of ELAVL1 sites towards the poly-A tail is specific for our dataset. The 3’UTR binding site distribution for ELAVL1 in HeLa contains a similar depletion towards the stop-codon but does not include the distinct enrichment of binding towards the poly-A tail. Per previous reports, the HeLa binding sites are equally distributed across the distal region of the 3’UTR (Lebedeva et al., 2011).

The most striking change observed between naïve and stimulated conditions was the increase in the percentage of exonic sites (from 26% to 48%), indicating a profound shift in binding-preference of ELAVL1 upon immune stimulation (Figure 1E and S1C). This divergence in the total number of sites and the change in binding site-specificity does not seem to be due to a subsampling issue as there were more unique and total reads per cluster in the stimulated condition samples (Wang et al., 2015).

From our PAR-CLIP data, we observed a major shift in not only the number of ELAVL1 binding sites between the naïve and stimulated states (~100,000 naïve to ~40,000 stimulated), but we also see a condition-specific difference in the proportion of sites that map to introns and the 3’UTR. The change in the distribution of ELAVL1 sites between the naïve and stimulated cellular states gives insight into the context-dependent mRNA-targeting of ELAVL1. From the RNA-Seq data, we observed that the mRNA levels (i.e., the potential ELAVL1 mRNA substrates) significantly change during an IRF3-driven immune response. Therefore, we wanted to understand how ELAVL1 mRNA targeting changes during a highly dynamic mRNA-substrate environment. Namely, we wanted to examine the proportion of ELAVL1 binding sites that are found on newly expressed stimulated specific transcripts. Furthermore, if ELAVL1 binds a target mRNA in both conditions, is the number and location of the ELAVL1 binding sites the same, and how do changes in binding site position affect the function of ELAVL1 on that transcript? We would predict that ELAVL1 would target upregulated transcripts in the 3’UTR to promote mRNA stabilization and gene expression.

### The mRNA target spectra of ELAVL1 differ in immune cell type

To investigate how ELAVL1 differentially targets mRNAs due to cellular condition, we first determined the conservation of ELAVL1 binding sites between the naïve and stimulated states and found that 27,323 clusters overlapped by at least one nucleotide (Figure 2A). This overlap comprises 27% of the sites in the naïve state and 70% of the sites in the stimulated state. At the mRNA transcript level, we found that the majority (5,051) of targets were bound in both naïve and stimulated conditions, though a notable number were uniquely found in the naïve (1,289) or stimulated (444) states (Figure 2B, Table S4).

**Figure 2.**
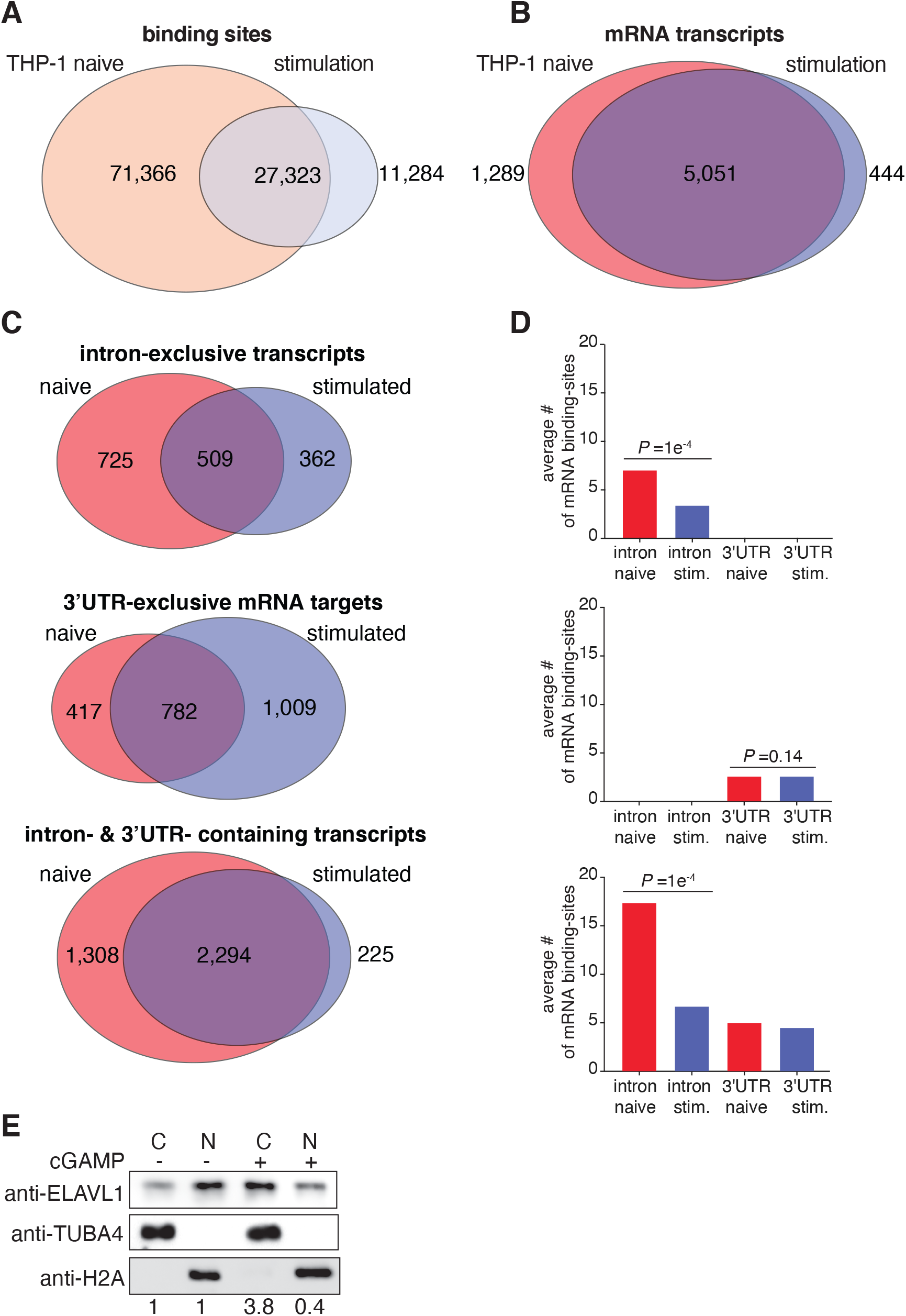
Innate immune stimulation pivots ELAVL1 binding towards 3’UTR sites. During immune stimulation, there is a significantly decreases in the total number of intronic binding sites transitioning the binding preference of ELAVL1 to the 3’UTR. (**A**) Venn diagrams showing the overlap of PAR-CLIP-defined ELAVL1 clusters between naïve and stimulated samples within mRNA or, (**B**), the overlap at the transcript level. (**C**) Venn diagrams indicating the number of transcripts in naïve and stimulated cells that are bound exclusively in the intron, 3’UTR, or both. (**D**) The average number of binding sites for the three mRNA location categories (intron-exclusive, 3’UTR-exclusive, or intron- and 3’UTR-containing transcripts) across conditions. (**E**) Immunoblot showing the cellular distribution of ELAVL1 (nuclear or cytoplasmic) during a naïve and stimulated cellular state. Tubulin (TUBA4) and histone 2A (H2A) are shown as localization controls. Quantitation of ELAVL1 bands are shown, relative to the corresponding compartment in the naïve state.

We wanted to parse this data further and investigate if the mRNAs that were shared, or uniquely bound, were differentially targeted by ELAVL1. Of the 6,340 mRNAs bound by ELAVL1 in the naïve state, > 94% of transcripts contained sites that mapped to intron-exclusive, 3’UTR-exclusive, or both; < 6% of transcripts contained sites within 5’UTR and/or CDS. A similar proportion was observed for the transcripts bound in the stimulated state. Consequently, we focused our analyses on transcripts that bore a distribution of intron and 3’UTR sites, which naturally divided into three populations: transcripts that contained ELAVL1 sites exclusively within: (1) introns, (2) the 3’UTR, or (3) both regions. In comparing the proportion and the absolute number of transcripts bound by ELAVL1 that were either intron- or 3’ UTR-exclusive, we observed a significant preference shift for the 3’UTR in the stimulated state. Conversely, the naïve state had nearly 50% more transcripts that were exclusively bound within introns (Figure 2C).

We next examined mRNAs that contained ELAVL1 binding sites within introns and the 3’UTR, as they represented the majority of ELAVL1 targets. Since these transcripts contained sites for both regions, we reasoned that a change in intra-transcript binding would give insight into how ELAVL1 targets RNA in a condition-specific manner. Previous work has shown that that the total number of ELAVL1 binding sites correlate with the extent of its regulation for that particular mRNA (Mukherjee et al., 2011). Therefore, we calculated the average number of ELAVL1 sites for each region, with the hypothesis that a change in intra-transcript binding preference would point to the region(s) of the mRNA that is relevant to its function during immune stimulation. In the stimulated state, we see a two-to three-fold reduction in the number of intronic binding sites for intron-exclusive transcripts and those that contained both intron and 3’UTR sites (Figure 2D). Among the mRNA targets that contained both intron and 3’UTR sites, the ratio of intron sites to 3’UTR sites shifted from 3:1 to 1.3:1 upon stimulation. Corroborating the observation that the changes are due to the loss of intronic binding sites, we found no significant change in the average number of 3’UTR sites upon immune stimulation. Altogether, our data show that immune stimulation leads to a loss of mRNA targets exclusively bound within introns, as well as the total number of intronic sites across all other transcripts – resulting in a net increase in the proportion of 3’UTR bound ELAVL1 targets. The change in intra-transcript target preference observed for ELAVL1 may be explained by its translocation to the cytoplasm as we and others have observed upon immune stimulation (Figure 2E) (Blanco et al., 2016; Grammatikakis et al., 2016; Lourou et al., 2019).

Given the condition-dependent binding site preference shift from intron to 3’UTR, we wanted to see if there was a change in motif usage by ELAVL1. Motif analysis (6-mer analysis) of the binding sites in both conditions located in 3’UTR and introns revealed a UUUUU and AUUUA rich RNA recognition element (RRE) (data not shown). Similar results were obtained from previous PAR-CLIPs from HeLa and HEK293, indicating that ELAVL1 does not change sequence-specific binding site preference in different cell types or conditions (Mukherjee et al., 2011). These data suggest that other factors, such as RNA secondary structure and competition with other RBPs, influence the ability of ELAVL1 to bind to its RNA targets.

### PAR-CLIP and RIP-Seq define enrichment criteria for ELAVL1 during an innate immune response

We assessed the quantitative changes in substrate binding by ELAVL1 as cells transitioned from the naïve to the immune activated state, given the rapid changes to the transcriptome that we observed. ELAVL1 functions through its targeting of AREs within mRNAs. By quantifying its association to targets, we can discern the salient binding properties that drive the relative enrichment differences caused by an IRF3-driven innate immune response. Therefore, we performed RIP-Seq for ELAVL1 (2017; Keene et al., 2006; Tenenbaum et al., 2000; Zhao et al., 2010; 2008). Flag-HA ELAVL1 from THP-1 cells was immunoprecipitated in the same manner as with PAR-CLIP but without UV crosslinking and RNA digestion. RNAs that co-immunoprecipitated with ELAVL1 were recovered and sequenced. A total of 3,459 and 3,406 PAR-CLIP identified mRNA targets in naïve and stimulated samples, respectively, were enriched over the IgG background by RIP-Seq (Figure 3A and 3B, Table S4). Overlapping the enriched targets from both conditions, we found that 2,114 mRNAs were enriched in a context-independent manner; 1,345 mRNAs were specific to the naïve state, while 1,292 were specific to the stimulated state (Figure S2A). We next discerned whether the enriched targets in the stimulated specific state represented just the upregulated mRNAs. Surprisingly, we found that a majority (61%) of the 1,292 transcripts were already expressed in the naïve state, and their expression values did not significantly change (< two-fold change) upon immune stimulation. The mean expression value for these existing transcripts was 10.7 CPM (all transcripts mean = 9.2 CPM). This data indicates that the differences in the target repertoire between the two cellular states are not solely due to changes in the expression levels of mRNAs. ELAVL1 competes for binding sites with other post-transcriptional regulatory elements including miRNAs or other RBPs, whose expression and activities are similarly dynamic and dependent on the cellular context (Dassi, 2017; Lu et al., 2014; Srikantan et al., 2012; L. E. Young et al., 2012).

**Figure 3.**
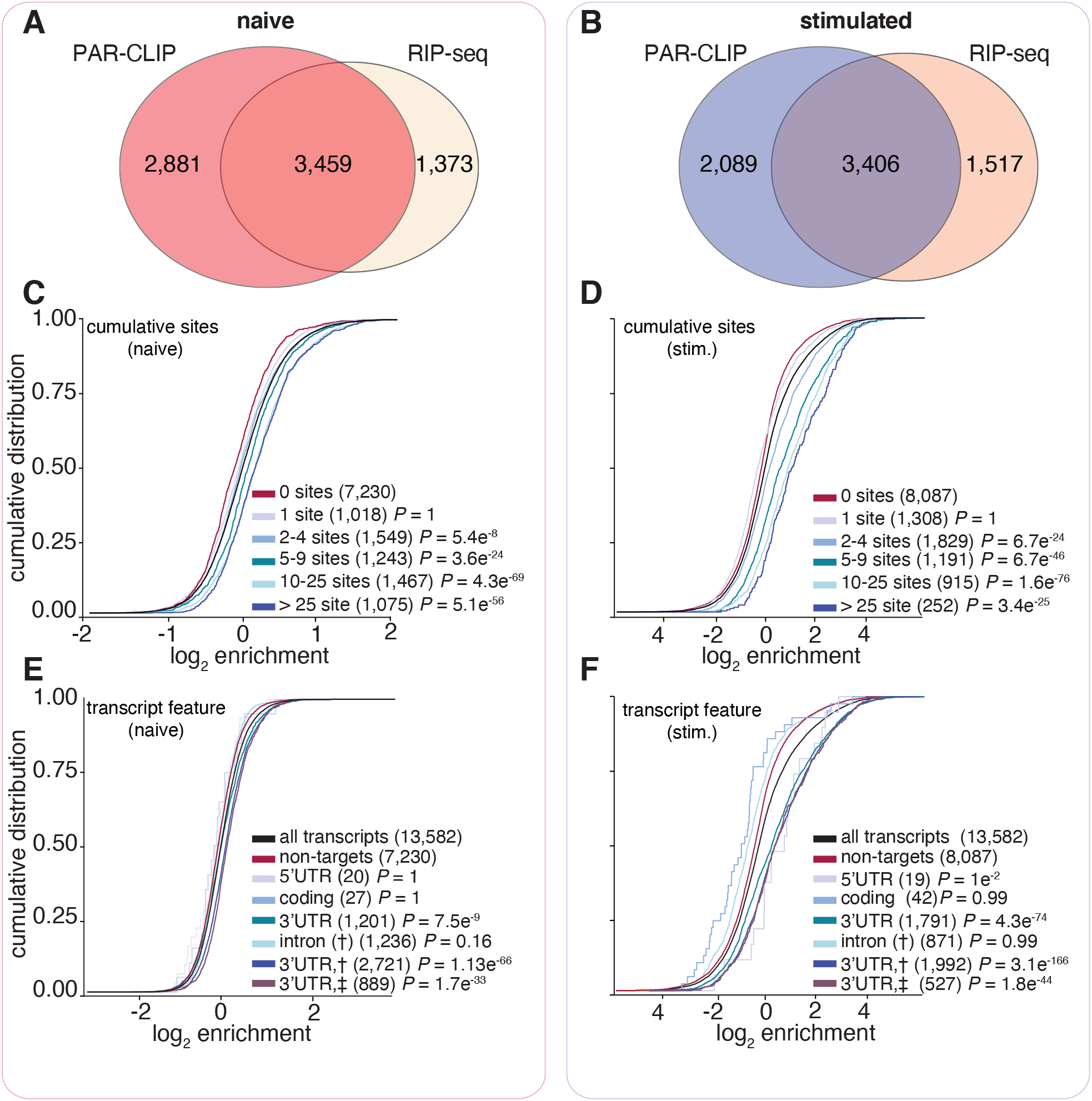
ELAVL1 RIP-Seq and PAR-CLIP define transcript enrichment criteria during immune stimulation. The total number of ELAVL1 binding sites confers greater enrichment, especially in the stimulated condition, and requires association at the 3’UTR. (**A**) Venn diagram of the overlap between mRNA transcripts defined as targets using both PAR-CLIP and RIP-Seq data in the naïve and (**B**) IRF3 stimulated state. (**C**) To test the hypothesis that an increasing number of ELAVL1 sites lead to transcript enrichment, we performed cumulative distribution fraction analyses for the mRNA targets of ELAVL1 in naïve or (**D**) stimulated conditions on the basis of the number of total binding sites indicated. (**E**) To test the hypothesis that the location of the binding site contributes to enrichment, we performed cumulative distribution fraction analyses for ELAVL1 mRNA targets in naïve or (**F**) stimulated cells. Transcripts were binned based on the indicated location of binding sites.

Reactome pathway analysis on target genes that were either naive-specific or shared showed that they encoded for proteins involved in transcriptional regulation by TP53 (R-HSA-3700989), processing of capped-mRNAs (R-HSA-72203), and cell cycle checkpoints (R-HSA-69620) (Figure S2B-C, Table S5) (Techasintana et al., 2015). These pathways comprise of more ubiquitous cellular processes that are generalizable across multiple cell types, often found as steady-state functions. By contrast, Reactome pathways enriched from targets specific to the stimulated state include toll-like receptor (TLR) signaling (R-HSA-168181), NF-kB (R-HSA-975183) and MAP kinase (MAPK) signaling (R-HSA-975138) (Figure S2D, Table 5). This observation is interesting because the majority of the targets that were enriched in the stimulated-specific state were expressed at similar levels in naïve cells, but not enriched - thus suggesting that ELAVL1 associates with transcripts belonging to immune signaling pathways regardless of mRNA levels (transcriptional output). Consistent with our observation, 71% (918/1,292) of these stimulated-specific enriched targets were also found as PAR-CLIP bound transcripts in a HEK293 study, yet only 122 were enriched; pathway analysis of these 122 did not yield enrichment in the TLR, NF-kB, or MAPK signaling terms (Mukherjee et al., 2011).

### 3’UTR binding determines the level of enrichment to context-dependent mRNA targets

To test the hypothesis that the frequency and position of ELAVL1 binding sites influence levels of enrichment in THP-1 cells, we examined the cumulative distribution of ELAVL1 target enrichment (RIP-Seq) based on PAR-CLIP binding site data. Independent of cellular state, transcripts with ≥ 2 binding sites showed significant enrichment compared to transcripts with one or zero sites. Furthermore, an increase in the number of ELAVL1 binding sites showed a positive correlation with enrichment (Figure 3C and 3D). Interestingly, we saw that enrichment levels were more pronounced in the stimulated state. Overall, for transcripts that had ≥ 2 sites in the stimulated state, there was nearly a 200% increase in fold-enrichment over non-targets, whereas targets (≥ two sites) in the naïve state were only nominally enriched (10%) over non-targets.

From our PAR-CLIP data, we observed a net decrease in the total number of binding sites per transcript in the stimulated state compared to naïve. However, we found that an increase in the number of sites per transcript correlated with greater enrichment – especially in the stimulated state. To reconcile these observations, we grouped transcripts based on the location of PAR-CLIP sites. We found that mRNA that contained at least one 3’UTR binding site had the highest levels of enrichment compared to binding sites in other transcript regions (Figure 3E and 3F). A majority of 3’UTR-bound transcripts also contained additional intronic sites, and previous reports showed that intronic binding contributed to enrichment (Lebedeva et al., 2011; Mukherjee et al., 2011). Therefore, we tested whether increasing numbers of intron bound sites led to greater enrichment but found no correlation (Figure 4A and 4B). Enrichment was dependent solely on the number of 3’UTR sites in both naïve and stimulated states. Of note, the 3’UTR-bound transcripts in the stimulated state were five times more enriched over non-3’UTR bound targets compared to the same populations in the naïve condition (Figure 4C-D). We also observed that the fractional occupancy of 3’UTR sites per transcript was significantly favored in the stimulated state (Figure 4E). This might explain the increase in the 3’UTR-specific enrichment in the stimulated state compared to the naïve.

**Figure 4.**
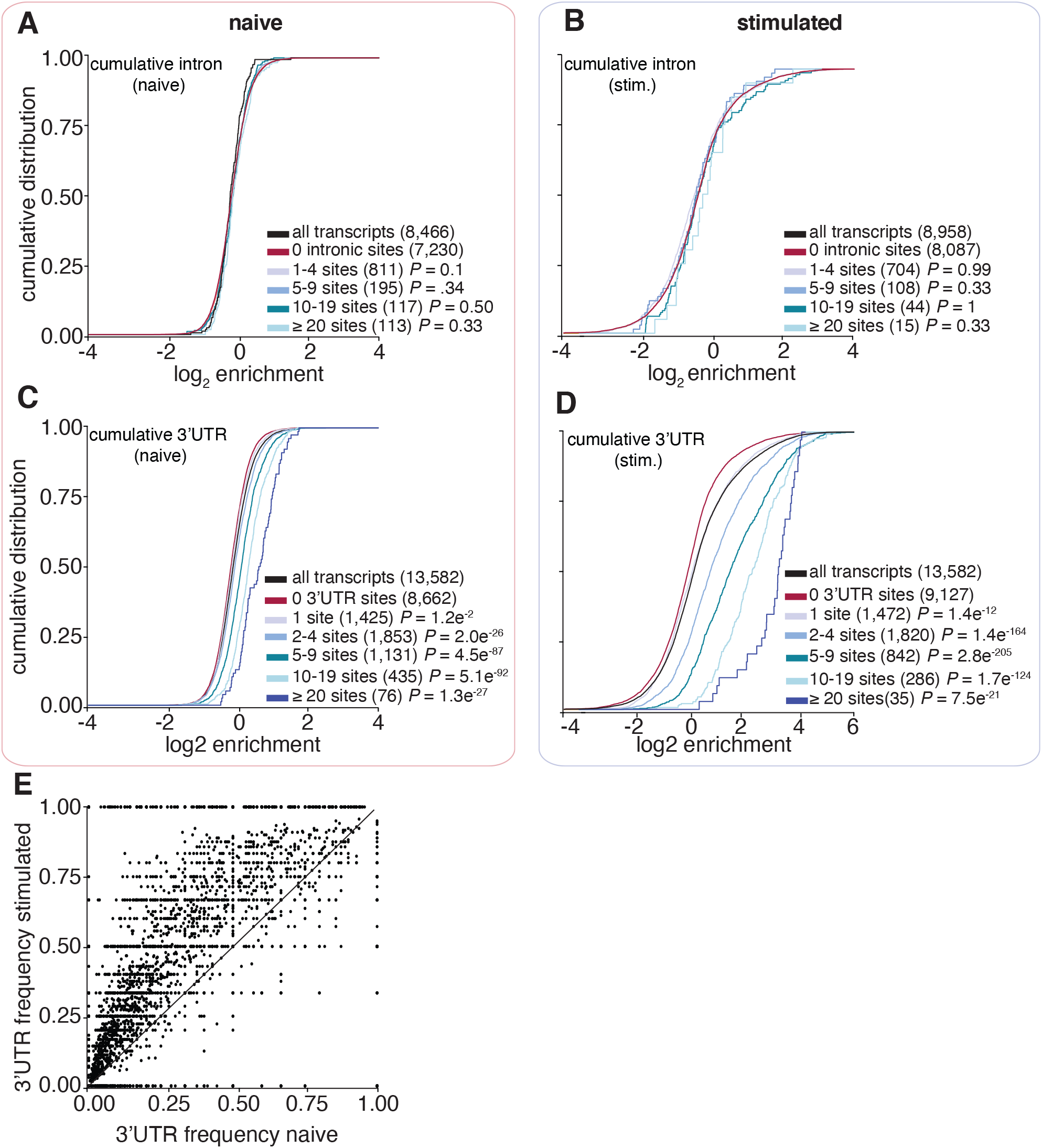
ELAVL1-mRNA enrichment is exclusively dependent on 3’UTR association and intensifies upon immune stimulation. Binding of ELAVL1 at 3’UTR sites drives the greatest levels, and is sufficient for, transcript enrichment when cells are immune activated. (**A, B**) Cumulative distribution function analyses were used to determine whether intronic binding versus 3’ UTR binding (**C**, **D**) confers greater transcript enrichment in naïve and stimulated states. (**E**) A scatterplot shows the bias of fractional occupancy, or frequency, of 3’UTR binding per given transcript for the stimulated state. The fraction of 3’UTR sites over the total number binding sites was plotted wherein the x-axis represents the naïve state and the y-axis indicates the stimulated state.

From the integration of RIP-Seq and PAR-CLIP datasets, we observed that an increasing number of 3’UTR sites correlates with greater enrichment with ELAVL1. Importantly, mRNAs bound by ELAVL1 in the immune-stimulated state showed significantly higher enrichment levels, demonstrating a stronger association and potentially indicating a more profound level of post-transcriptional gene regulation. However, the binding and enrichment of a transcript with an RBP is not directly equivalent to its regulatory fate, given that any single mRNA is potentially subject to other post-transcriptional factors that may impose a stronger regulatory effect. In most reported cases, the primary function of ELAVL1 is through the stabilization of its target transcripts (Herdy et al., 2015; Srikantan et al., 2012; Takeuchi, 2015b; Turner and Díaz-Muñoz, 2018). ELAVL1 competes over AREs on transcripts, opposing negative post-transcriptional regulators like RNA-induced silencing machinery or ZFP36 (TTP), which can recruit the CCR4-NOT1 deadenylase complex (Fu and Blackshear, 2016). Consequently, to best understand the functional impact of our enriched ELAVL1 targets, we utilized SLAM-Seq to quantitatively measure transcript decay rate (Herzog et al., 2017).

### ELAVL1 stabilizes a subset of 3’UTR targets involved in innate immune signaling

By overlapping SLAM-Seq results with our RIP-Seq and PAR-CLIP datasets, we were able to precisely identify the consequences of ELAVL1 absence on the stabilities of its target transcripts during an innate immune response (Figure 5A-B). Slam-Seq uses 4SU to metabolically label nascent RNA transcripts, which is subsequently chased with unlabeled uridine. Thiol-alkylation of RNA generates chemical adducts that induce reverse transcriptase-dependent deoxycytosine substitutions at 4SU positions during cDNA library preparation (T-to-C substitutions). The ratio of unlabeled to T-to-C containing reads across all the timepoints is used to calculate RNA half-life for each expressed transcript (Herzog et al., 2017; Neumann et al., 2019). Here, we performed SLAM-Seq during an IRF3-driven innate immune response. Parental and ELAVL1-KO THP-1 cells were labeled with 4SU for 16 h, followed by washout and chase using uridine. Cells were collected at various time points after the chase, and extracted RNAs were processed for high-throughput sequencing. We determined the half-lives of nearly ~5,000 transcripts that were shared across parental and ELAVL1-KO datasets and found their median half-lives to be 6.7 hours and 6.4 hours, respectively (Figure 5C and Table S6), indicating comparable global RNA stability independent of condition type. Importantly, the difference in the median half-lives between ELAVL1 targets and non-targets is greater when ELAVL1 is present (1.8 h), in comparison to the median half-life difference when it is knocked out (0.7 h) (Figure 5D). These data indicate that the half-lives of ELAVL1 target transcripts are more similar to non-targets upon its knockout.

**Figure 5.**
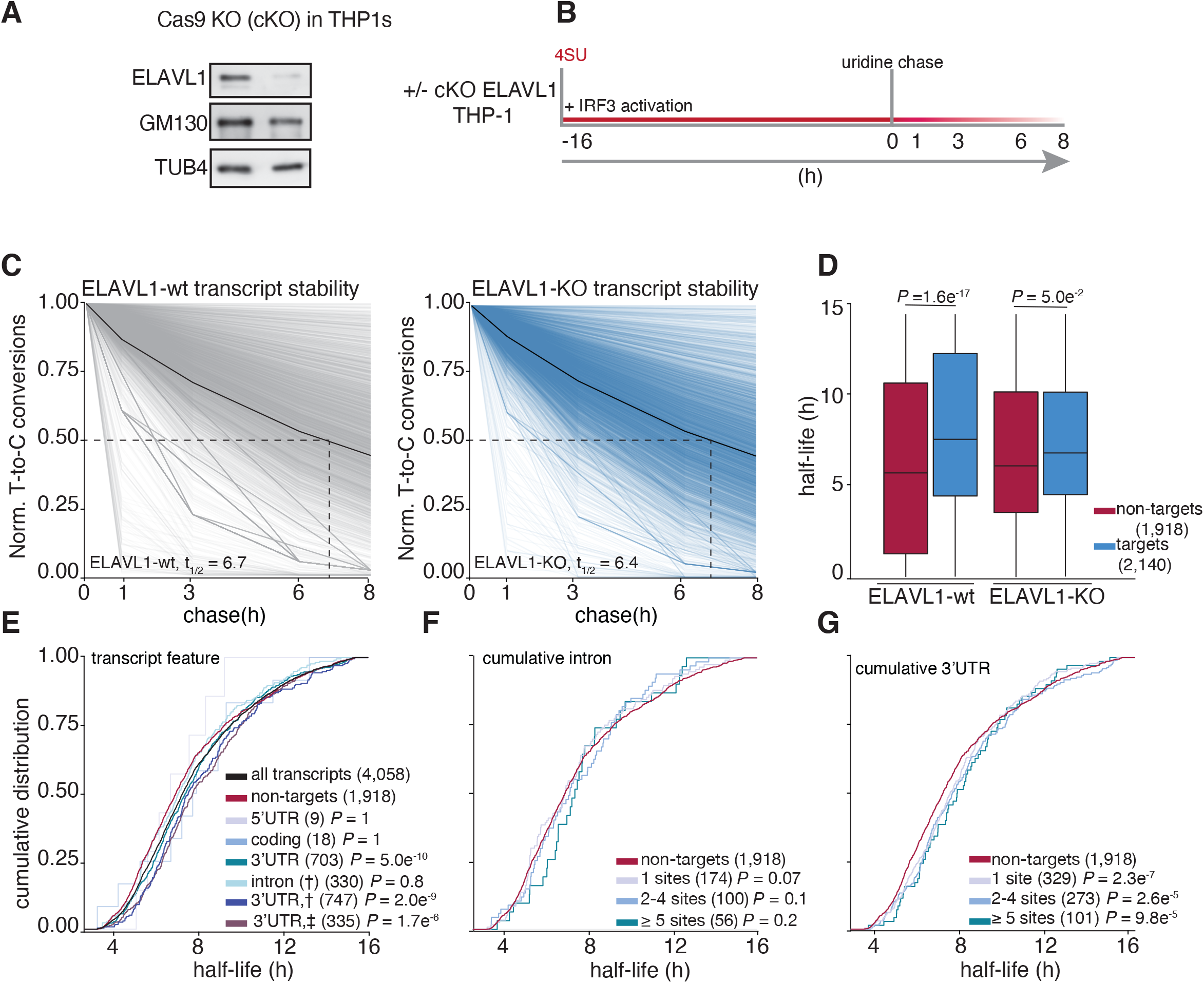
Analysis of transcriptome-wide mRNA stability in the absence of ELAVL1. Overlapping SLAM-Seq transcript stability data with RIP-Seq enrichment and PAR-CLIP binding site information defines which target transcripts are most affected by the loss of ELAVL1. (**A**) Immunoblot staining for the presence of endogenous ELAVL1-wt and ELAVL1-KO THP-1 cells. (**B**) Schematic of the SLAM-Seq experiment setup. THP-1 cells (+/−KO) were stimulated and 4SU labeled for 16 hours before wash and uridine chase. Timepoints for SLAM-Seq were 0, 1, 3, 6, and 8 hours after uridine wash. (**C**) The decay of T-to-C conversions after the uridine chase were determined by fitting the data to a single-exponential decay model to derive mRNA half-lives (dotted-line). Graphs show that the RNA stabilities over time, for each measured transcript in the ELAVL1-wt and ELAVL1-KO cells, are overall similar. Solid black line indicates the median T-to-C conversion rate over time; dotted black line denotes median half-life (t_1/2_), as indicated per condition. (**D**) Box plots showing the median half-lives of non-targets and targets in the ELAVL1-wt and ELAVL1-KO cells. (**F**) To test if location of ELAVL1 binding sites contributes to increased RNA stability, we performed cumulative distribution fraction analyses of the RNA half-lives measured grouping transcripts of the basis of binding sites location. (**G**). Cumulative distribution fraction analyses were used to determine whether the RNA half-lives of targets are affected by the number of intronic- or (**H**) 3’UTR-binding sites. Data not shown for CDS and 5’UTR targets because of too few targets.

In the parental THP-1 cells, we observed that transcripts with at least one 3’UTR binding site had statistically significant longer RNA half-lives (t_1/2_ = 7.5 h) than non-targets (t_1/2_ = 5.7 h); whereas, 5’UTR, coding-, or intronic-bound targets were not statistically significant (Figure 5E). We noticed that transcripts that have binding sites either in the intron and 3’UTR (t_1/2_ = 7.6 h) or intron and 3’UTR plus another location (t_1/2_ = 7.5 h) tended to have slightly longer RNA half-lives than transcripts bound exclusively in the 3’UTR (t_1/2_ = 7.4 h). Similar to our analysis of transcript enrichment, of comparing the intron- or 3’UTR-exclusively bound transcripts, we found that only the presence and increase in the number of 3’UTR sites conferred greater stability (Figure 5F and 5G). Interestingly, targets with >10 3’UTR binding sites exhibited half-lives 3 hours longer than non-targets (t_1/2_ = 8.9 h versus 5.7 h). In looking at the change in target transcript half-lives between KO and wt, no feature other than 3’UTR conferred any significant stability effect (Figure S3A).

To assess whether target transcripts that were enriched based on 3’UTR content were specifically stabilized by ELAVL1, we examined their behaviors when ELAVL1 was knocked out. The half-lives of only the enriched 3’UTR-containing ELAVL1 transcripts were significantly reduced in the absence of ELAVL1 (Figure 6A). The half-lives of the non-enriched transcripts bound by ELAVL1 were not as affected by the absence of ELAVL1, suggesting that the decay rates of these transcripts are either partially or completely independent of ELAVL1 regulation, despite being bound. Moreover, transcripts with increasing number of ELAVL1 sites have the greatest decrease in half-lives further supporting this 3’UTR stability signature (Figure 6B). Among the most 3’UTR enriched targets (n=1,141), we noted that a substantial fraction (50%) are ISGs, and collectively they had a greater change in half-life in the knockout, compared to enriched transcripts that are not ISGs (Figure 6C). ISGs such as *NFKB1*, *IRF9*, *IFIT5*, and *CXCL11*, are less stable in the absence of ELAVL1, in contrast to ISGs for which we had zero evidence of ELAVL1 association or enrichment (Figure 6D).

**Figure 6.**
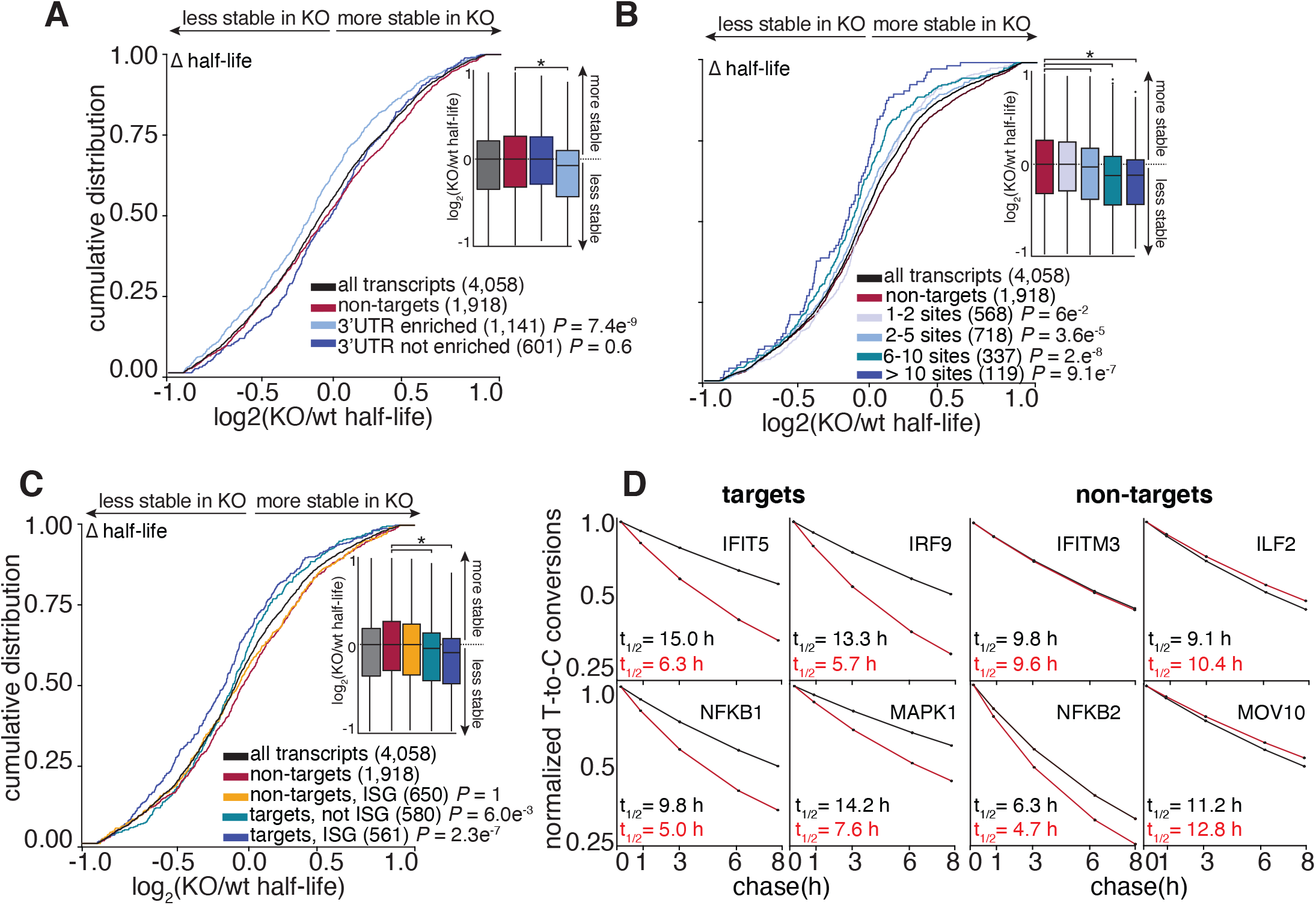
The RNA half-lives of highly enriched ELAVL1 target ISGs are the most affected by its loss. The RNA stabilities of enriched transcripts with an increasing number of 3’UTR sites were the most affected by the absence of ELAVL1 compared to non-targets and targets that were not enriched. (**A, B**) To test if enrichment predicted a decrease in transcript half-life in the ELAVL1-KO, we performed cumulative distribution analyses of the log_2_ fold-change in half-life (KO/wt), binning transcripts based on enrichment. Insets show a box plot of the log_2_ fold-change (KO/wt) in half-life based on indicated groupings. (**C**) To test if transcripts classified as ISGs have a greater decrease in half-life in the KO compared to non-ISG targets, we plotted cumulative distribution function graphs of the log2 fold-change in half-life (KO/wt). (**D**) Transcript stability plots and calculated RNA half-lives of the indicated ISG mRNA targets or non-targets are shown; ELAVL1-wt (black line) or ELAVL1-KO THP-1 cells (red line)

Pathway analysis showed that the ELAVL1-regulated targets (n = 409), which we defined as 3’UTR enriched targets whose half-life decreased by 1.5 fold-change in the absence of ELAVL1, encode for proteins associated with endocytosis, transcriptional dysregulation in cancer, and multiple innate immune signaling pathways (Figure 7A and Table S7). Of the immune-relevant pathways, ELAVL1 regulated components of interleukin-17 (R-HSA-448424), TNF-pathway (R-HSA-75893), and Toll-like receptor signaling (R-HSA-168164). Other signaling pathways, such as MAPK (R-HSA-450294), and apoptosis (R-HSA-109606) terms, were also enriched. MAPK signaling, along with NF-kB and IRFs, are important for generating an immunoreactive state in the presence of a pathogen (Arthur and Ley, 2013). In the case of TLR signaling, ELAVL1 binds directly and stabilizes the adaptor protein (TRAF6), a kinase involved in integrating upstream PRR activity (TAK1), and the transcription factor itself (NF-kB) – which are all positive regulators of the pathway. Overall, we found that these innate immune pathways form a network containing regulators of ISG expression (STAT3, MAPK, IRF9, FOS, NF-kB, RIPK2) (Figure 7B) (Gilchrist et al., 2012; Mostafavi et al., 2016). Thus, our data indicate that ELAVL1 stabilizes the transcripts of ISGs and ISG regulators critical for a cell to mount an immunoreactive state.

**Figure 7.**
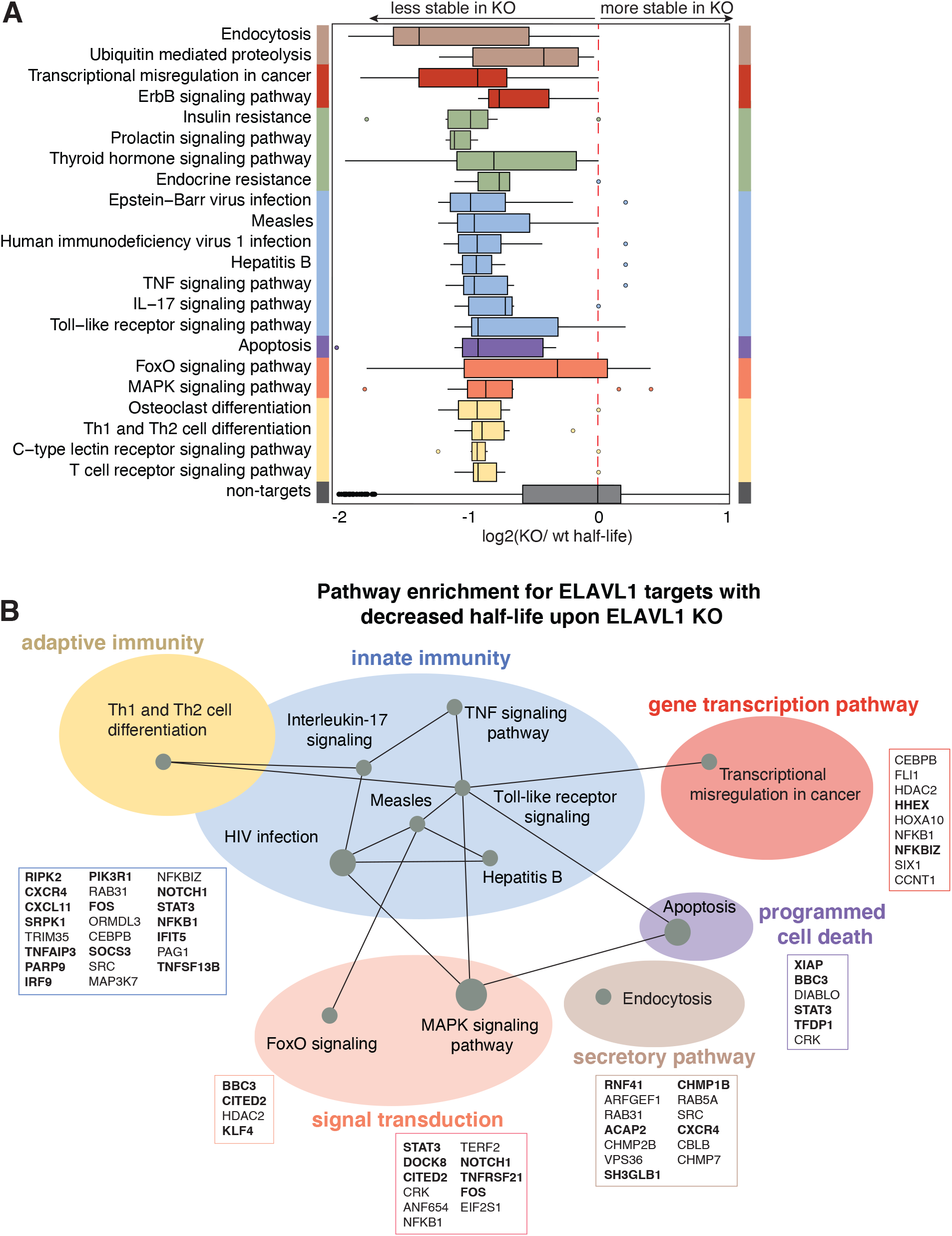
Canalization of ELAVL1 function towards the post-transcriptional regulation of immunologic pathways by IRF3 stimulation. (**A**) Box plot grouping transcripts based on the top enriched KEGG and Reactome pathway terms of the ELAVL-regulated targets protein components. Pathways were determined based on using clusterProfiler R/Bioconductor package. Fisher’s exact test using Benjamini-Hochberg FDR < 0.05. (**B**) Pathways of functional targets of ELAVL1. Each node represents a specific pathway, and colored circles represent closely related pathways. Connections represent shared genes between pathways. Boxes show specific mRNA transcripts and ISGs (bolded) in each pathway that are targeted by ELAVL1.

Of note, we found that a surprising number of the highly regulated transcripts were already bound by ELAVL1 in the THP-1 naïve state. However, their enrichment level (based on rank) were significantly higher upon immune stimulation. This increase in enrichment was concomitant with higher 3’UTR fractional occupancy - reinforcing the importance of a transition to 3’UTR binding. Only ~10% of our ELAVL1-regulated targets were found as enriched or even affected by the knockdown of ELAVL1 in HEK293 cells – underscoring the importance of examining ELAVL1 targets under changing transcriptomic contexts.

## DISCUSSION

We present a multi-layered analysis of high-throughput transcriptomics of the targeting and functional outcomes of the RBP ELAVL1 during an innate immune response. Using expression and PAR-CLIP data, we identified 98,689 naïve and 38,607 stimulated ELAVL1 RNA-binding sites. To date, this is the first report comparing how the binding properties and the targeting of an RBP change between steady-state and innate immune conditions. We find that ELAVL1 largely transitions to binding the 3’UTRs of mRNA transcripts during an innate immune response. 3’UTR binding is an absolute prerequisite for enrichment, and knockout of ELAVL1 led to widespread destabilization of its enriched transcripts. Specifically, we found that highly regulated targets had a three-fold average reduction in their stabilities, losing 30 to 80% of their original half-lives. Importantly, ELAVL1-regulated targets encode for ISGs and their transcriptional regulators, suggesting that ELAVL1 contributes at multiple levels of a pro-inflammatory response.

Two other ELAVL1 binding site reports performed in HeLa and BMDMs, show that ELAVL1 mostly binds the 3’UTR of mRNA targets (Lebedeva et al., 2011; Sedlyarov et al., 2016). In both of these reports, investigators immunoprecipitated ELAVL1 using endogenous antibodies for PAR-CLIP. In our hands, we found that endogenous antibodies obstructed RNA-protein interactions. Therefore, we used anti-Flag antibodies in this study and compared our results to the previous PAR-CLIP work performed in HEK293 cells which used the same antibodies. Accordingly, we also observed a high proportion of ELAVL1 binding occurring at intronic sites, particularly during the naïve state. Nonetheless, our data did not show a link between ELAVL1 intronic sites and an increase in mature mRNA stability (Lebedeva et al., 2011; Mukherjee et al., 2011). Those previous reports used microarrays that could detect introns and pre-mRNAs to discover an intron-dependent role of ELAVL1 for stabilizing pre-mRNAs. Whereas our approach with SLAM-Seq was geared towards measuring the RNA decay rates of mature transcripts. While intuitively, changes in pre-mRNA levels would be predicted to contribute to the eventual levels of mature transcripts, we did not observe an intron-dependent effect on mature mRNAs. It is conceivable that our approach was insufficiently sensitive to detect intronic-dependent stability effects or that the two ELAVL1-dependent mechanisms are partially distinct.

One of the strongest signatures we were able to identify was a shift in the binding preferences by ELAVL1 for the 3’UTRs of mRNAs. We interpret the dominance of the 3’ UTR preference in defining enrichment and stability, upon immune stimulation of THP-1 cells, is likely due to subcellular re-localization of ELAVL1 to the cytoplasm – where it would be able to bind the 3’UTRs of mature mRNAs, free of introns. As previously reported, ELAVL1 can be post-translationally modified by numerous kinases and methylases, ultimately dictating its nucleocytoplasmic localization and RNA-binding preference (Grammatikakis et al., 2016). Future studies using subcellular-restricted forms of ELAVL1 (e.g., phospho-ablated or phospho-mimetic) could add to the granularity of our understanding of how ELAVL1 transitions from binding intronic versus 3’UTR sites and its movement out of the nucleus. However, the subcellular localization of ELAVL1 can only partially explain our work, since signal transduction and associated transcriptomic differences between naïve and stimulated states also play a role in defining the ELAVL1-regulated mRNA (Abe et al., 2012; Rabani et al., 2011). Given the presence of other post-transcriptional regulatory factors and elements, ELAVL1 likely competes with the actions of other RBPs, miRNA silencing machinery, and possibly viral RNAs that could act as competitors or sponges to titrate its activities away from cellular targets (Barnhart et al., 2013; Dassi, 2017; Hentze et al., 2018; Kim et al., 2020; Lu et al., 2014). Therefore, when characterizing the function of a given RBP and its biological role, it is important to consider how cellular context-specific factors can have a profound impact on how a target enriches with an RBP and the extent of its regulation.

Since transient associations can be identified between potential RNA targets and RBPs by CLIP methodologies, there is a need to quantify enrichment levels as a measure of interaction affinity. The stoichiometry of interactions between the limited number of RBP molecules and its changing RNA substrate pool makes it difficult to predict enrichment and post-transcriptional regulatory impact using information gleaned strictly from steady-state data, particularly for RBPs with a preference for highly redundant recognition sequences (Ascano et al., 2011). High throughput discovery methods, such as RNAcompete and RNA bind-n-Seq, show that most RBPs tested have convergent RNA recognition sequences (Dominguez et al., 2018; Lambert et al., 2014; Ray et al., 2017). For 27 different RBPs (including ELAVL1), the preferred 6-mer binding site overlapped with the top-ranked 6-mer site of at least one other RBP, despite having distinct RNA-binding domains (Dominguez et al., 2018). This observation underscores that many RBPs have similar and short binding-motif preferences of low-complexity (Adinolfi et al., 2019; Nussbacher and Yeo, 2018). Though these studies are valuable in discovering the primary and flanking sequence preferences of RBPs, these methods often analyze a single RBP in isolation and do not include competitor RBPs and miRNAs that will influence the RNA-substrate structure or the availability of a particular RNA binding site. Therefore, relying on sequence motifs and predicted RNA structure is insufficient to delineate the specific binding sites and the functional RNA targets of RBPs in specific cells and contexts. In recognizing these limitations, we integrated multiple high-throughput sequencing datasets to differentiate between RNA transcripts that are simply ‘sampled’ versus *bona fide* targets that are regulated during an immune signaling event.

For our functional analysis, we focused on the majority of targets, which exhibited a reduction in their half-lives in the absence of ELAVL1. We termed these mRNAs ELAVL1-regulated transcripts and performed pathway analysis to understand the cellular pathways that ELAVL1 could regulate during an immune response. Many of them (*NFKB1*, *IRF9*, *TAK1*, *STAT3*, *MAPK1*, and‡‡ *MAPK*9) encode protein components involved in innate immune signaling that positively drive the expression of ISGs. These genes reinforce upstream signaling transduction events triggered by pathogen and nucleic-acid sensing, leading to enhanced cytokine and interleukin production, and cellular differentiation. We speculate that ELAVL1 regulates these central signaling components to sensitize the cell for dealing with a pathogenic infection – allowing the cell to quickly integrate incoming pathogen-or damage-associated molecular pattern triggered signaling. Furthermore, our data support the idea of an “RNA regulon” where an RBP can coordinate the expression of a group of functionally-related mRNAs (Keene, 2007; Simone and Keene, 2013). ELAVL1 regulates the mRNA of several biological processes within immune signal transduction pathways-from adaptor protein (TRAF6) to transcription factor (NF-kB, IRF9). ELAVL1 also regulates components of endocytosis, which plays a role in cytokine signaling and TLR receptor trafficking (Kurgonaite et al., 2015; Lund and DeLotto, 2014). Importantly, these transcripts were strongly regulated by ELAVL1 upon immune stimulation, despite many of these targets already being sampled by the RBP under naïve states.

With the increasing interest in understanding the function of RBPs in gene regulation, numerous laboratories have undertaken essential and broad surveys of the target spectra of RBPs (Castello et al., 2016; Darnell, 2010; Hafner et al., 2010; König et al., 2010; Nostrand et al., 2020; Ule et al., 2003; Van Nostrand et al., 2016). But it is important to recognize that RBPs represent a broad class of gene regulators that are post-translationally modified and are sensitive to cellular signaling events and changing transcriptomes. In examining the contribution of ELAVL1 to immune stimulation, we provide a general framework for studying other RBPs subject to analogous changes to a dynamic transcriptome or signal transduction event. This is especially relevant during host-pathogen interactions when substrate RNAs compete for post-transcriptional gene regulation by cellular proteins, leading to dramatic changes in the balance of host versus pathogenic transcript binding with limited trans-acting factors.

## ACKNOWLEDGEMENTS

We would like to thank Kelly Barnett for computational help in the analysis of the PAR-CLIP and RIP-seq datasets; Monica Bomber for technical help with generation of the ELAVL1-KO; and the Vanderbilt Technologies for Advanced Genomics (VANTAGE) core for sequencing. Finally, we would like to thank members of the Ascano laboratory for their support, collegiality, and critical review of the manuscript. This work was supported by the National Institutes of Health 1R35GM119569-01 (M.A.), Vanderbilt University Dept. Biochemistry start-up funds (M.A.), the Chemistry-Biology Interface training grant 5T32GM065086-14 (K.R.), the Chemical Biology of Infectious Disease training grant 5T32AI11254-02 (S.A.), and Clinical and Translation Science Award No. UL1 TR002243 (K.R.).

## AUTHOR CONTRIBUTIONS

Conceptualization and Methodology, K.R. and M.A.; Software and Data Curation, K.R. and S.A.; Formal Analysis, K.R., Investigation, K.R., S.A., and C.R.; Resources, K.R., B.K., S.A., and N.M.; Writing - Original Draft, K.R.; Writing – Review and Editing, K.R., S.A., M.A., and N.M.; Visualization, K.R.; Funding Acquisition, M.A. and K.R.; Supervision, M.A.

## STAR METHODS

### RESOURCE AVAILABILITY

#### Lead Contact

Manuel Ascano

#### Materials Availability

Further information and requests for resources and reagents should be directed to and will be fulfilled by the Lead Contact, Manuel Ascano. All plasmids and stable cell lines generated in this study are available without restrictions from the Lead Contact and/or through Addgene.

#### Data and Code Availability

All code used for sequencing analysis and figure generation is accessible at https://github.com/Ascano-Lab.

### EXPERIMENTAL MODEL AND SUBJECT DETAILS

#### Cell lines and culture

Human THP-1s monocytes (male) were cultured in RPMI (Gibco) with 10% fetal bovin serum (FBS from Peal Serum), 100 μg/ml streptomycin (Gibco), 100 U/ml penicillin (Gibco), 25 μg/ml blasticidin and 100 μg/ml hygromycin (Invivogen).

#### Plasmid construction

For the cloning of the lentiviral expression construct, the ELAVL1 coding sequence was PCR amplified from THP-1 cDNA, introducing *attB*-sites for the Flip-IN-recombinase system. The PCR product was gel purified and then recombined into the pLenti-CMVtight-Flag-HA-DEST-Blast plasmid. The Flag-HA-tag lentiviral inducible expression vector pLenti CMVtight Blast Flag-HA-DEST was constructed by insertion of Flag-HA-tag from pFRT_TO_DEST Flag-HA (#26361, Addgene) into the plasmid pLenti CMVtight Blast DEST (w762-1) (#26434, Addgene).

#### Lentiviral Production and generation of Inducible expressing Flag-HA ELAVL1

For lentiviral production, HEK293T were first cultured in 15 cm plates in high glucose DMEM (Gibco) supplemented with 10% FBS. HEK293T were then transfected using Lipofectamine 2000 (Invitrogen) according to manufacture suggestion with 9 μgs of lentiviral vector and 9 μg viral particle packaging vector, 6.75 μg psPAX2 (12260, Addgene) and 2.25 μg pMD2.G(12259, Addgene). 48 hours after transfection, the viral particle-containing supernatant was collected and spun-down at 3000 g for 15 mins. The supernatant was then concentrated and purified by layering the supernatant over a 20% sucrose cushion in TNE buffer (50 mM Tris-HCl [pH 7.2], 0.1 M NaCl, and 1 mM EDTA) and ultra-centrifuged at 25,000 g for 4 hours in a Beckman SW32Ti rotor. Viral pellets were then resuspended in fresh DMEM media and filtered through 0.45 μm syringe filter unit (Millex-HV). For viral transduction, THP-1-rtTA cells were spun-inoculated (800 g for 2 hours at 32°C) with an MOI ~100. Two days after viral inoculation, cells were moved into selection media 25 μgs/ml blasticidin and 100 μg/ml hygromycin. Expression of ELAVL1 was then verified via immunoblot using both ELAVL1 endogenous antibody or anti-HA antibody.

#### RNA-Sequencing and Library Prep

RNA from 1 × 10^6^ THP-1s were collected at indicated timepoint and were washed with 1 x PBS. For stimulated samples, we activated cells for 16 hours with the EC50 of encapsulated cGAMP (Shae et al., 2019). Cells were then resuspended in 1 mL of TRizol. RNA was extracted following the manufacture’s protocol. Total RNA was converted into cDNA and sequenced using NEBNext DNA Library Prep Kit for Illumina on the Illumina NovaSeq6000 platform using PE150 at the Vanderbilt Technologies for Advanced Genomics (VUMC VANTAGE). Fastq files were pre-processed with trim-galore with the default settings (http://www.bioinformatics.babraham.ac.uk/projects/trim_galore/) to remove any adapter contamination and then aligned to the human genome (Genocode, hg19) with STAR mapper (Dobin et al., 2012). DeSeq2 (Love et al., 2014) was used to calculate differential expressed genes.

#### PAR-CLIP

PAR-CLIP was performed as previously described (Garzia et al., 2016; Hafner et al., 2010) with minor adjustments. In brief, 3-5 × 10^9^ THP-1 cells were doxycycline induced and labeled with 100 μM 4SU 24 hours before harvesting and UV_365nm_ irradiation. Stimulated THP-1 cells were additionally treated with 70 nM (EC50) cyclic GMP-AMP 16 hours before harvest. After crosslinking, THP-1s were lysed using NP-40 lysis buffer (50 mM HEPES [pH 7.5], 150 mM KCl, 2 mM EDTA, 1 mM NaF, 2% (v/v) NP40, 0.5 mM DTT, Roche EDTA-free protease inhibitor) and incubated with Dyna-protein G beads (Invitrogen) coupled with anti-FLAG M2 antibody (Sigma) for ~ 2 hours at 4°C. Beads were washed with high-salt buffer and then underwent CIP, and T4 PNK mediated 5’-end RNA radiolabeling with [γ-^32^P]-ATP. Flag-tagged ELAVL1 crosslinked to RNA was then was resolved on a 4-20% Bis-Tris, NuPage gradient gel (Invitrogen). The band corresponding to ELAVL1 protein was cut out. The protein:RNA complex was then electroeluted out of the gel and treated with proteinase K (Roche). RNA was then size selected and underwent both 3’ (MultiplexDX Inc.) and 5’ adapter (Illumina compatible) ligation and was reverse transcribed into cDNA. cDNA library was sequence on the NextSeq Illumina platform at Hudson Alpha.

#### Defining binding sites

PARalyzer (Corcoran et al., 2011; Mukherjee et al., 2014) was used to define crosslinking sites from the PAR-CLIP data. PARalyzer calculates the T-to-C fraction which serves as a quality index that is calculated based on the frequency of a given uracil (thymidine) to be substituted with a cytosine. For groups of reads (> 5 unique reads), kernel density estimates were calculated for both reads with and without T-to-C conversions. Clusters (i.e., binding sites) were defined as group of transcripts that had a higher kernel density for T-to-C converted reads over unmodified reads. https://github.com/ohlerlab/PARpipe

#### Motif Analysis

For the 6-mer analysis, we counted the most frequent 6-mers from each unique PAR-pipe called clusters annotated as either intron or 3’UTR using BioStrings R/Bioconductor (Pages et al., 2020). To calculate the top enriched 6-mers for ELAVL1, we regressed the 6-mer frequency relative to a reference library of annotated 3’UTRs or introns (Mukherjee et al., 2018).

#### RIP-Sequencing

Immunoprecipitation was performed as previously described in the PAR-CLIP method section without UV crosslinking or RNAse treatment. An anti-IgG immunoprecipitation (IP) was used to subtract background RNA expression that are intrinsic of IPs. Following anti-FLAG and anti-IgG IP, beads were added to 1 ml TRizol (Ambion) and RNA was extracted following the manufacture’s protocol and total RNA was submitted to Vantage sequencing core.

#### Statistical Analysis

The significance difference between cumulative distribution fractions were calculate using Wilcoxon.Test, and *P*.values were adjusted using the Bonferroni method to the number of observations. Student’s t test was calculated to compare the means of different categories (e.g., difference in the number of binding sites between two conditions).

#### Reactome and KEGG analysis

Reactome and KEGG analysis was performed using R/Bioconductor packages clusterProfiler with default settings. (Yu et al., 2012).

#### RT-qPCR

RNA collected and extracted using Trizol (Ambion). Concentration of total RNA was determined using NanoDrop 2000 (ThermoFisher). Equal amounts of total RNA for each sample were reverse transcribed using SuperScript III (ThermoFisher) with randon hexamer. Real-time PCR reactions were done with FastSYBR Green Plus Master Mix (Applied Biosystems) and a StepOnePlus qPCR machine (Applied Biosystems). Target Ct values were normalized to *TUBA1A* Ct values and used to calculate ΔCt. Relative mRNA expression of target genes was then calculated using the ΔΔCt method (2^ΔΔCt^).

#### Nucleo-cytoplasmic Fractionation

Harvested cells were washed with 1 x PBS and resuspended in hypotonic lysis buffer (10 mM HEPES, 60 mM KCl, 1 mM EDTA, 0.075% (v/v) NP40, 1 mM DTT, and 1 mM EDTA-free Roche PMSF) and incubated on ice for 10 minutes. Cell lysis was then centrifuge at 4°C (15,000 rcfs, 5 minutes) to pellet the nuclear fraction. Top cytoplasmic fraction was removed and place into a new tube. Nuclear fraction pellet was washed twice in hypotonic lysis buffer without NP40 and then lysed with hypertonic lysis buffer (20 mM Tris Cl, 420 mM NaCl, 1.5 MgCl_2_, 0.2 mM EDTA, 1 mM EDTA-free PMSF (Roche) and 25% (v/v) glycerol).

#### Antibodies and immunoblotting

The antibodies to ELAVL1 (anti-ELAVL1, ab170193), anti-TUBA4A (ab7291), from Abcam; anti-Histone 2A (JBW301) is from Sigma. anti-GM130 (12480) is from Cell Signaling. Samples were separated by SDS-PAGE. After electrophoresis, proteins were semi-dry transferred (Bio-Rad) to nitrocellulose membranes (Hybond-ECL, GE Life Science). Protein membranes were processed via a standard immunoblot protocol followed by enhanced chemiluminescent detection (Luminata Forte ECL, Millipore) using a chemiluminescence imaging system (ChemiDoc MP, Bio-Rad).

#### Generations of Cas9 sgRNA knockout in THP-1 monocytes

crRNAs were designed using the CRISPR design tool in Benchling [Biology Software] (2019) retrieved from https://www.benchling.com/crispr/ and ordered from IDT. To assemble Cas9/sgRNA RBPs, first Alt-R crRNAs and trcRNAs were first reconstituted in Nuclease Free Duplex buffer (IDT). An equimolar ratio of crRNA and trcRNA were added to a nuclease-free tube and denatured by heating at 95 °C for 5 minutes. The oligo duplex was then cooled at room temperature for 10 minutes prior to adding Cas 9 nuclease enzyme (IDT) to assemble RNPs (Vakulskas et al., 2018). Duplexed oligos and Cas9 were then incubated at room temperature for 20 minutes prior to THP-1 electroporation.

For electroporation, 2 × 10^6^ THP-1 cells per sample were counted and washed twice with 1 x PBS. Assembled Cas9 RNPs and washed cells were then suspended in 100 μl of R buffer (NeonTransfection) and electroporated in 100 μl tip with the NeonTransfection unit (1600V, 10 ms, 2 pulses). Electroporate cells were then added to pre-warmed THP-1 media (RPMI + 10% FBS) without antibiotics and cultured in an incubator 37 C + 5% CO_2_ for 48 hours before changing the media.

#### SLAM-Seq

THP-1 (ELAVL1-wt and ELAVL1-KO) cells were seeded the day before the experiment at a density of 2 × 10^6^ cells/ ml. For the pulse, cells were labeled with 100 μM 4SU and stimulated with 70 nM (EC50) cyclic GMP-AMP 16 hours before harvest. For the chase, cells were washed twice in 1x PBS and then incubated with RPMI + 10% FBS supplemented with 10 mM of uridine (Sigma). Cells were harvested, washed in 1x PBS, and added to TRIzol at the respective timepoints (0, 1, 3, 6 and 8 hours after chase). Total RNA (~5 μgs per sample) was extracted and then treated with 10 mM iodoacetamide (sigma) as described in Herzog et .al, 2017 in *Protocol Exchange* (DOI: 10.1038/protex.2017.105). SLAM-Seq libraries were prepared using the Lexogen QuantSeq 3’ mRNA-Seq Library Prep Kit FWD for Illumina (Cat. No. 015.24) following the manufacturer’s instructions. Libraries were sequenced using Illumina NextSeq 550 in a 75 bp single-end mode. SLAM-Seq libraries were analyzed as previously described in (Herzog et al., 2017) and (Neumann et al., 2019). Briefly, read converged normalized T-to-C conversion rates were generated using the SLAM-DUNK pipeline.

To calculate RNA half-lives, T-to-C conversion rates were normalized to chase onset (0-hour timepoint) and used to fit a first-order decay reaction in R using the min-pack. lm package (Elzhov et al., 2016).

