## Supplemental Figures and Legends for "ELAVL1 Primarily Couples mRNA Stability with the 3’UTRs of Interferon Stimulated Genes"

### SUPPLEMENTAL FIGURE LEGENDS

**Supplemental Figure 1: PAR-CLIP identifies and maps the cell-type specific and condition specific binding sites of ELAVL1**
(**S1A**) Gene expression analysis using qPCR measuring the mRNA levels of *IFNB1*. THP-1 cells were stimulated with the STING agonist cGAMP (EC50) and RNA was collected at indicated timepoints.

(**SIB**) Bar graph showing the number of clusters that mapped either to exons and introns across the two cellular conditions. Inset shows the average number of unique reads for each exonic- or intronic- cluster across the conditions.

(**S1C**) Venn diagram showing the overlap of target mRNAs from HEK293 (Mukherjee et al. 2011) and THP-1 naïve.

(**S1D**) Reactome pathway analysis for the mRNAs that were unique bound by THP-1s compared to HEK293.

(**SIE**) Metagene analysis showing the normalized distribution of binding sites across introns or (**SIF**) the 3'UTR across both conditions.

(**SIG**) Metagene analysis showing the distribution of binding sites in the 3'UTR that are 200 nucleotides downstream of the poly-A tail.

**Supplemental Figure 2: PAR-CLIP and RIP-Seq identify enriched transcripts that are condition independent or dependent**
(**S2A**) Venn diagram showing the overlap between the bound (PAR-CLIP) and enriched (RIP-Seq) mRNA transcripts across conditions.

(**S2B**) Bar graph of a Reactome pathway analysis for each group of transcripts that were either shared (from S2A) between the two conditions or unique bound and enriched in the naïve (**S2C**) or (**S2D**) the stimulated states.

**Supplemental Figure 3: KO determines transcripts whose half-lives are dependent on the presence of ELAVL1**
(**S3A**) Cumulative distribution plot of the $\log_2$ foldchange (KO/wt) in half-life showing the group of transcripts whose half-lives are most affected by the loss of ELAVL1.

**Supplemental Figure 1** **Rothamel et al., 2020**

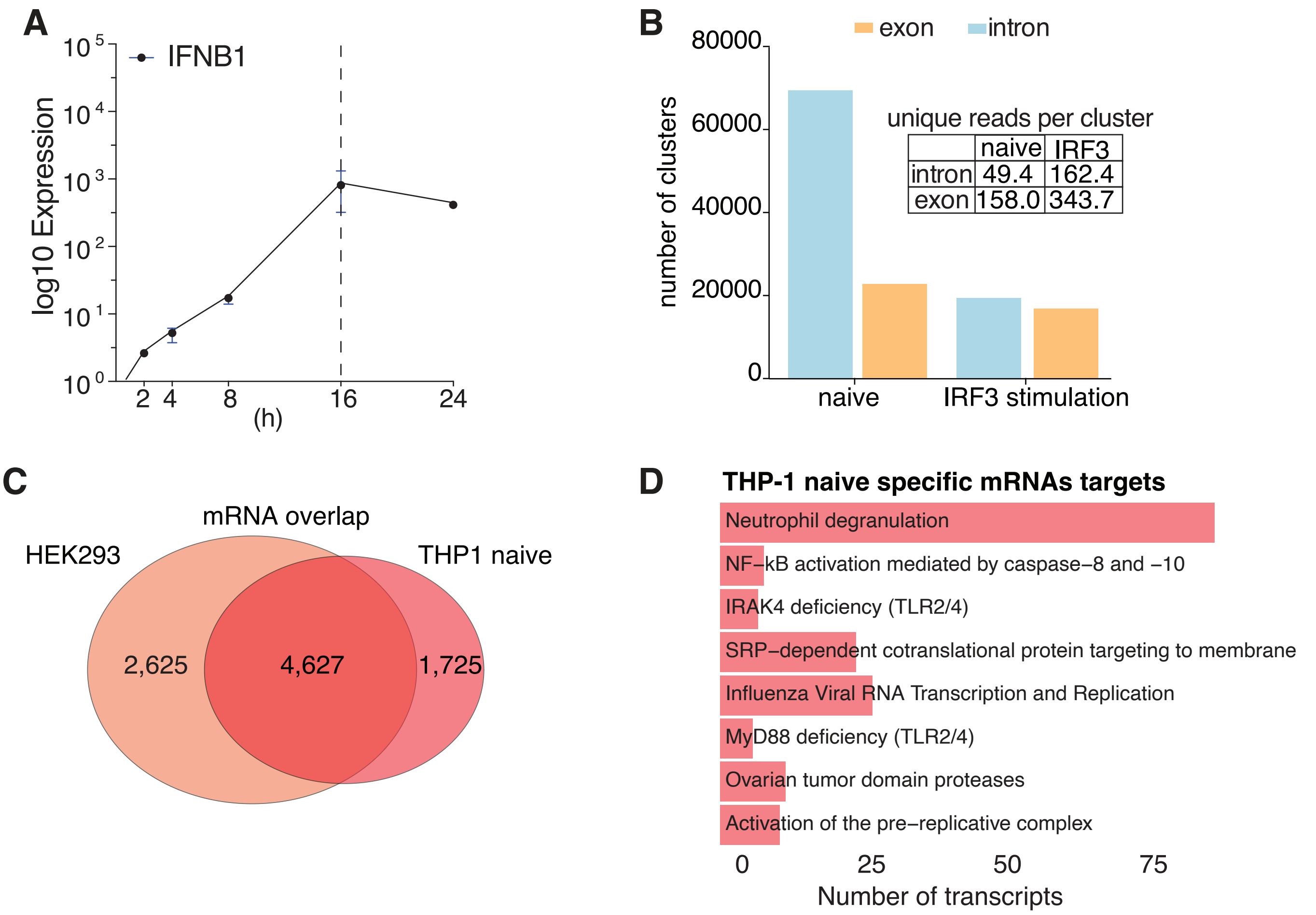

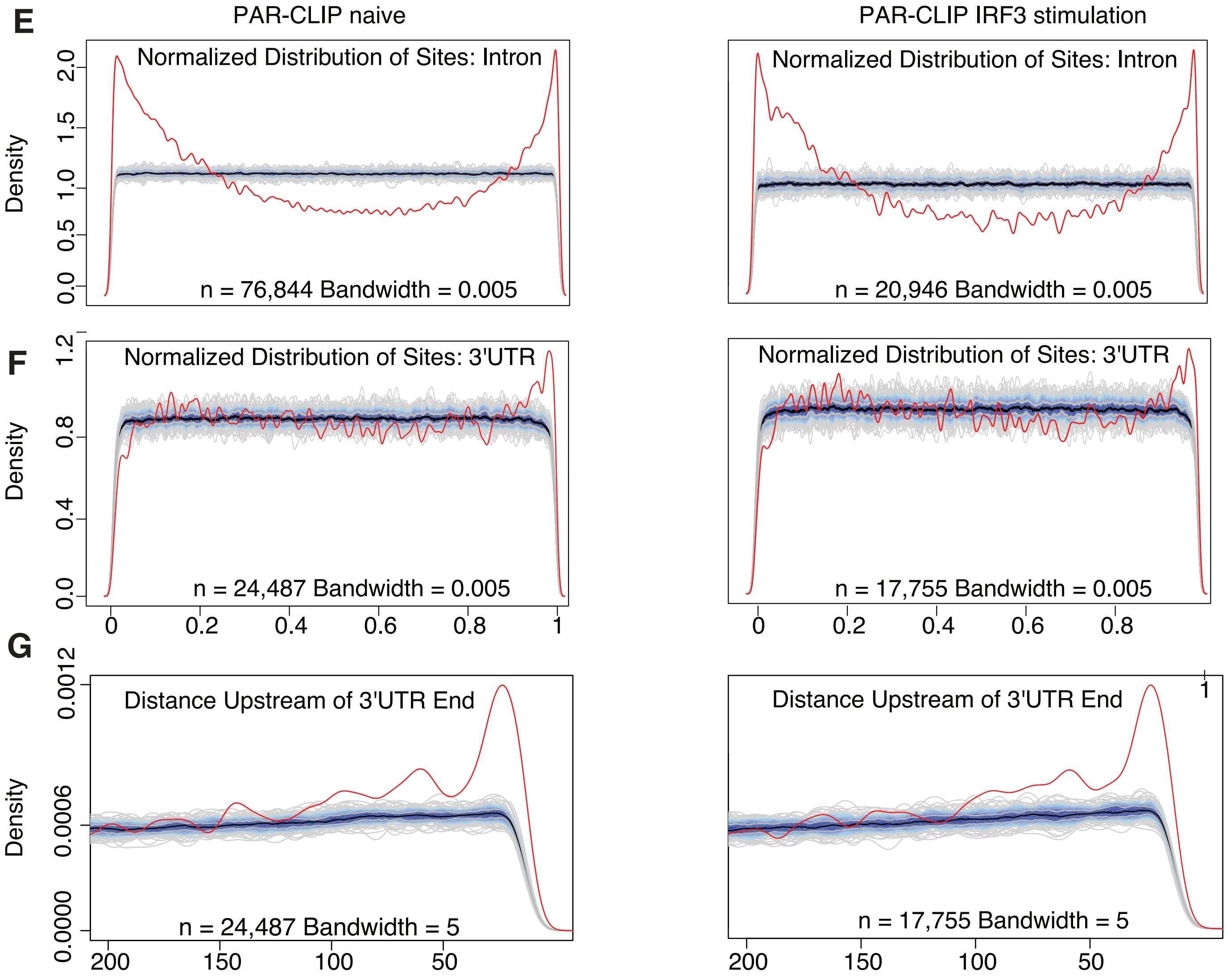
E
PAR-CLIP naive
PAR-CLIP IRF3 stimulation
Normalized Distribution of Sites: Intron
Density
n = 76,844 Bandwidth = 0.005
n = 20,946 Bandwidth = 0.005
F
Normalized Distribution of Sites: 3'UTR
n = 24,487 Bandwidth = 0.005
n = 17,755 Bandwidth = 0.005
G
Distance Upstream of 3'UTR End
n = 24,487 Bandwidth = 5
n = 17,755 Bandwidth = 5

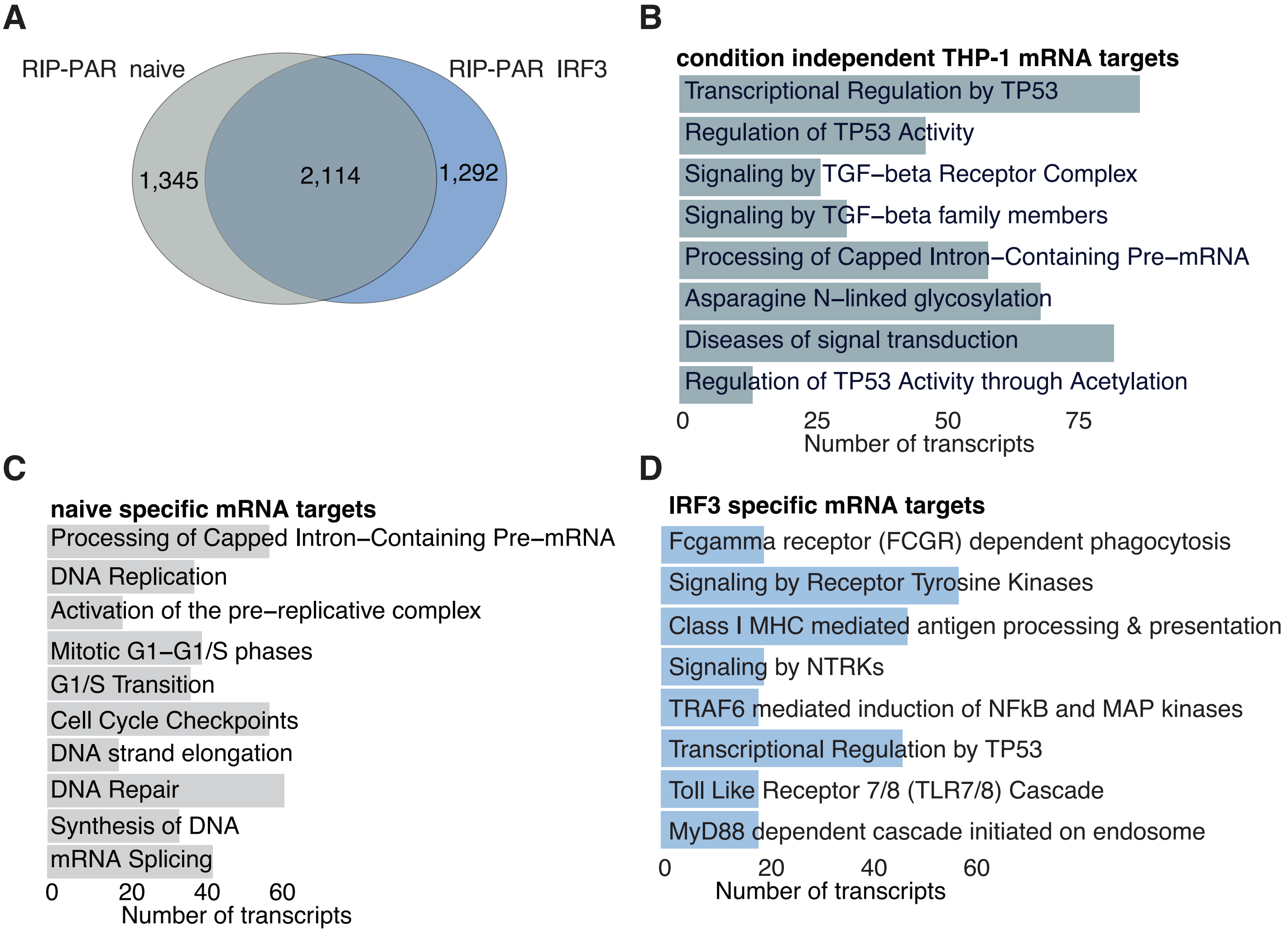
A
RIP-PAR naive
RIP-PAR IRF3
1,345
2,114
1,292
B
condition independent THP-1 mRNA targets
Transcriptional Regulation by TP53
Regulation of TP53 Activity
Signaling by TGF−beta Receptor Complex
Signaling by TGF−beta family members
Processing of Capped Intron−Containing Pre−mRNA
Asparagine N−linked glycosylation
Diseases of signal transduction
Regulation of TP53 Activity through Acetylation
0
25
50
75
Number of transcripts
C
naive specific mRNA targets
Processing of Capped Intron−Containing Pre−mRNA
DNA Replication
Activation of the pre−replicative complex
Mitotic G1−G1/S phases
G1/S Transition
Cell Cycle Checkpoints
DNA strand elongation
DNA Repair
Synthesis of DNA
mRNA Splicing
0
20
40
60
Number of transcripts
D
IRF3 specific mRNA targets
Fcgamma receptor (FCGR) dependent phagocytosis
Signaling by Receptor Tyrosine Kinases
Class I MHC mediated antigen processing & presentation
Signaling by NTRKs
TRAF6 mediated induction of NFkB and MAP kinases
Transcriptional Regulation by TP53
Toll Like Receptor 7/8 (TLR7/8) Cascade
MyD88 dependent cascade initiated on endosome
0
20
40
60
Number of transcripts

### Supplemental Figure 3

**A**

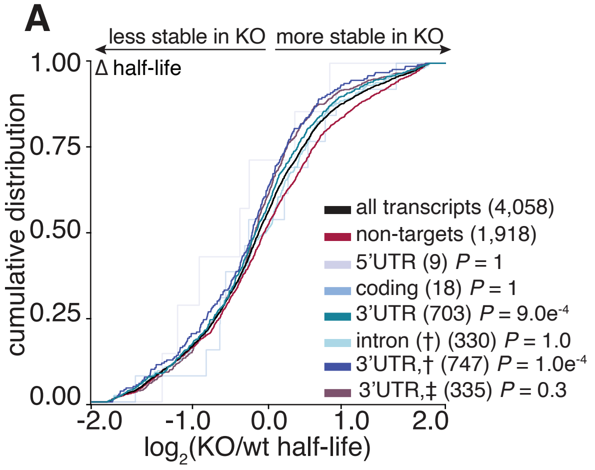
